# Transitory enhancement of GATA2 chromatin engagement during early erythroid differentiation

**DOI:** 10.64898/2026.03.05.709895

**Authors:** John W Hobbs, Samuel J. Taylor, Rajni Kumari, Nayem Haque, Lou Lou Victor, Ulrich Steidl, Robert A. Coleman

## Abstract

Erythroid differentiation requires precise regulation of transcription factor binding to chromatin targets as hematopoietic progenitors relinquish multipotency and activate lineage programs. GATA2 maintains progenitor identity and is thought to be progressively silenced as GATA1 levels rise; however, the precise changes in GATA2 chromatin binding kinetics during this transition remain undefined. Here, we combined live-cell single-molecule imaging in cell lines and primary mouse progenitors with CUT&Tag chromatin profiling to define GATA2 activity during erythropoiesis. Single-molecule tracking resolved two interaction modes: short-lived (<1 s) searching interactions and long-lived (>5 s) binding. Surprisingly, early erythroid differentiation was characterized by a transitory strengthening of long-lived GATA2 chromatin engagement. This manifested as increased residence time of GATA2 bound to chromatin in G1E-ER4 cells and an expansion of the long-lived bound population in HPC7 cells and primary mouse progenitors. This transitory phase of enhanced engagement declined upon further differentiation. Genome-wide mapping identified regulatory elements selectively occupied by GATA2 during this early transition state, revealing promoter-proximal sites enriched for GATA/RUNX motifs and distal elements containing composite GATA/E-box signatures. Together, our imaging and chromatin profiling indicate that GATA2 chromatin engagement is kinetically remodeled at the onset of differentiation, with early recruitment targets partitioning into distinct promoter- and enhancer-associated subclasses. These results support a model in which transcription factor kinetics constitute a dynamic chromatin engagement layer that characterizes the GATA2-to-GATA1 transition.

## Introduction

Cell-fate decisions during differentiation are orchestrated by transcription factors (TFs) that coordinate gene expression programs in a cell-type- and context-dependent manner (Lambert et al. 2018; Levine et al., 2014). In hematopoiesis, the commitment of multipotent progenitors requires transcriptional mechanisms that activate lineage-specific programs while restricting multipotency (Laurenti and Göttgens, 2018; Orkin and Zon, 2008). Hematopoietic TFs accomplish these functions by binding cis-regulatory elements and assembling complexes with cofactors that modulate transcriptional output (Heinz et al. 2015).

GATA2 is essential for maintaining hematopoietic stem and progenitor cell (HSPC) identity (Yamamoto et al. 1990; Katsumura and Bresnick 2017; de Pater et al. 2013). As a zinc-finger DNA-binding protein, GATA2 supports HSPC proliferation, survival, and the transcriptional programs that sustain progenitor potential (Tsai et al. 1994; Tsai and Orkin 1997). During erythropoiesis, *Gata2* expression decreases as *Gata1* expression increases. Although distinct in function, both factors recognize the WGATAR consensus motif. This property enables a regulatory transition in which the GATA2 protein promotes *Gata1* transcription with the GATA1 protein subsequently replacing GATA2 at shared genomic binding sites. This “GATA switch” is central to erythropoiesis (Johnson et al. 2020; Ross et al. 2012; Weiss et al. 1997). Disruption of this regulatory axis results in bone marrow failure, immune dysfunction, and a predisposition to MDS and AML (Homan et al. 2021; Spinner et al. 2014; Wlodarski, Collin, and Horwitz 2017).

Several model systems have been used to study GATA2 function during erythroid differentiation. The G1E-ER4 cell line represents an important model for dissecting the GATA1-induced regulatory switch (Weiss et al. 1997). G1E cells, derived from *Gata1*-null fetal liver, enable direct analysis of the GATA switch because re-expression of GATA1 induces terminal erythroid maturation via displacement of GATA2 from chromatin (Fujiwara et al. 2009; Suzuki et al. 2013). The widely used G1E-ER4 derivative cell line expresses a GATA1–estrogen receptor ligand binding domain fusion (GATA1-ER), allowing synchronized erythroid induction through controlled nuclear entry of GATA1. In contrast, Hematopoietic Progenitor Cell-7 (HPC7) cells maintain endogenous GATA2 and GATA1 regulation and represent a cytokine-responsive progenitor state in which GATA2 cooperates with factors such as FLI1, LMO2, SCL/TAL1, and RUNX1 to sustain multipotency (May et al. 2013; Tijssen et al. 2011). Mouse reporter models, including GATA2VENUS, have further defined *Gata2* expression dynamics in vivo (Ahmed et al. 2020). Although they have clarified many aspects of the GATA2-to-GATA1 transition, these systems rely primarily on static or population-level measurements, leaving the real-time dynamics of chromatin engagement unresolved.

Transcription factor interactions with chromatin are highly dynamic. Single-molecule imaging studies in mouse ES cells and cancer cell lines have shown that TFs alternate between brief chromatin interactions and multi-second long-lived engagements (Gebhardt et al. 2013; Liu et al. 2014; Morisaki et al. 2016). It is unknown how transcription factor chromatin binding kinetics change during differentiation. Specifically in hematopoiesis, researchers have not directly measured whether GATA2 chromatin engagement changes in magnitude or duration during the initiation of erythroid differentiation in live progenitors.

The necessity of resolving these dynamics at the single-cell level is underscored by recent studies revealing substantial heterogeneity and stochasticity in transcription states of hematopoietic cells. Single-molecule imaging of mRNA demonstrated that lineage-associated transcription factors, including *Gata1*, *Gata2*, and *Spi1*, are expressed at low copy number and frequently co-expressed within individual progenitor cells, despite appearing mutually exclusive in population-averaged or sequencing-based analyses (Wheat et al. 2020). Single-cell proteomic studies further showed that lineage commitment is accompanied by gradual, quantitative remodeling of transcription factor protein abundance rather than discrete binary switches (Palii et al. 2019). Together, these findings suggest that hematopoietic cell states are dynamic and probabilistic, and that transcription factor abundance alone may be insufficient to capture the regulatory mechanisms that govern lineage entry.

Here, we use a combination of single-molecule TF protein imaging and CUT&Tag chromatin profiling to define how GATA2 chromatin interaction modes and genomic occupancy change during erythroid differentiation. Together, these complementary approaches establish a dynamic framework revealing that erythroid entry is defined by a previously unappreciated distinct kinetic transition state.

## Results

### Single-molecule imaging reveals discrete kinetic modes of GATA2 chromatin binding in hematopoietic progenitor cells

To visualize GATA2 behavior at single-molecule resolution, we immobilized live hematopoietic progenitors on CD43-coated glass dishes (Loeffler et al. 2018) for highly inclined laminated optical sheet (HiLo) microscopy. We expressed GATA2 as a HaloTag (Los et al. 2008) fusion (GATA2–Halo) containing a destabilization domain, allowing temporal control of protein levels via ligand-dependent stabilization. Upon addition of Shield-1, the fusion protein was stabilized and labeled with limiting Janelia Fluor-646 HaloTag ligand to achieve the sparse density required for single-molecule tracking. Shield-1–dependent accumulation of GATA2-Halo was confirmed in live HPC7 cells (Figure 1A; Supplementary Video 1)

**Figure 1.**
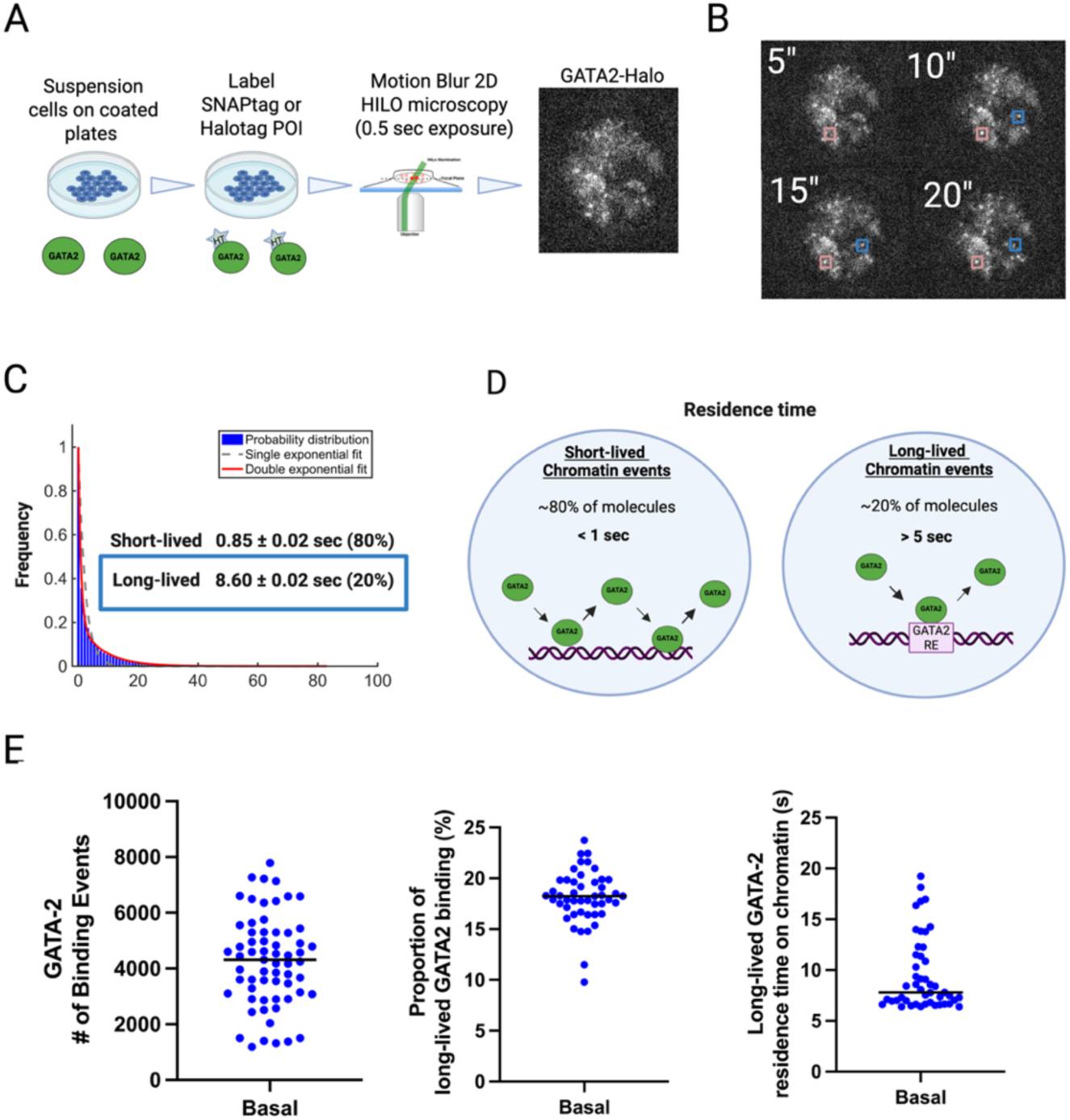
Single-molecule imaging reveals discrete GATA2 kinetic modes in hematopoietic progenitor cells. **(A)** Schematic of the live-cell single-molecule imaging workflow. Suspension cells were secured to the surface of an imaging dish using CD43-mediated immobilization and imaged by HiLo microscopy using 500-ms exposure times to preferentially detect chromatin-bound GATA2 molecules with multi-second residence times. **(B)** Representative time-lapse sequences illustrating two measurable GATA2 interaction modes. Short-lived interactions (<1 s) appear as brief sampling contacts in which GATA2 rapidly binds and unbinds chromatin. Long-lived interactions (>5 s) correspond to multi-second chromatin-bound events in which GATA2 remains localized at a single site. **(C)** Residence time distribution of GATA2 trajectories from a single representative cell fit with a double-exponential model, resolving a short-lived kinetic component (<1 s) and a long-lived component (>5 s) corresponding to chromatin engagement. **(D)** Schematic depiction of the interaction framework used throughout this study. Left: short-lived target-search behavior, in which GATA2 rapidly engages and disengages chromatin during chromatin scanning. Right: long-lived chromatin engagement, in which GATA2 remains localized at a defined regulatory element (RE) for several seconds. **(E)** Quantification of baseline kinetic parameters in **Basal progenitor cells (0 h)**, including total binding events per cell, fraction of molecules in the long-lived binding population (>5 s), and per-cell dwell-time distributions. Each point represents a single cell (N = 47), revealing substantial cell-to-cell heterogeneity in GATA2–Halo chromatin binding. Created with BioRender Hobbs, J. (2026) https://biorender.com/ux0rzr9

Single-molecule time-lapse imaging with a 500 ms exposure blurred out freely diffusing molecules, leaving only chromatin-bound GATA2–Halo visible as distinct fluorescent spots (Figure 1B; Supplementary Video 2). Trajectories were reconstructed and analyzed using the STRAP pipeline (Haque and Coleman 2025). The resulting dwell-time distribution was best fit by a double-exponential model, resolving short-lived (<1 s, ∼80%) and long-lived (>5 s, ∼20%) binding GATA2-Halo molecules (Figure 1C). Consistent with established single-molecule frameworks (Gebhardt et al. 2013; Liu et al. 2014), we attribute the short-lived population to non-specific chromatin scanning (Figure 1D, left) and the long-lived population to stable, sequence-specific regulatory engagement (Figure 1D, right).

Our data analysis pipeline provides several metrics related to chromatin-binding kinetics at the single-cell level, including (i) the total number of binding events, (ii) the proportion of molecules in the long-lived binding population (>5 s), and (iii) long-lived residence-time distributions. Plotting these metrics across a population of basal progenitor cells revealed substantial cell-to-cell heterogeneity of GATA2–Halo chromatin binding (Figure 1E). This heterogeneity suggests that individual cells within an otherwise defined progenitor population exhibit markedly different levels of GATA2 engagement with chromatin. For subsequent cell-state comparisons, we therefore focused primarily on the long-lived binding fraction, which reflects chromatin engagement associated with regulatory function (Figure 1D). Short-lived (<1 s) binding events were quantified separately and are reported in Figure Supplement 1.

### Transitory enhancement of GATA2 chromatin engagement during early erythroid differentiation in G1E-ER4 cells

The G1E-ER4 cell line provides a well-established model for synchronized erythroid differentiation through inducible nuclear entry of GATA1 (Weiss et al. 1997; Fujiwara et al. 2009; Suzuki et al. 2013). In this system, treatment with 4-hydroxytamoxifen (4-OHT) drives nuclear entry of GATA1–ER, initiating erythroid differentiation. Single-molecule imaging was performed in cells undergoing Basal progenitor (0 h), Early erythroid (2 h), and Late erythroid (24 h) states (Figure 2A; Supplementary Video 2). We confirmed expected erythroid progression using orthogonal protein and chromatin-based assays (Figure Supplement 2A-C). Western blot analysis showed progressive accumulation of GATA1 protein across the time course, while genome-wide CUT&Tag profiling revealed reciprocal changes in GATA factor occupancy at canonical erythroid regulatory loci, including *Kit*, *Gata2*, and *Klf1*. At these GATA-switch genes, GATA2 occupancy decreased while GATA1 binding increased during differentiation, consistent with prior studies (Suzuki et al. 2013) and validating our staging.

**Figure 2.**
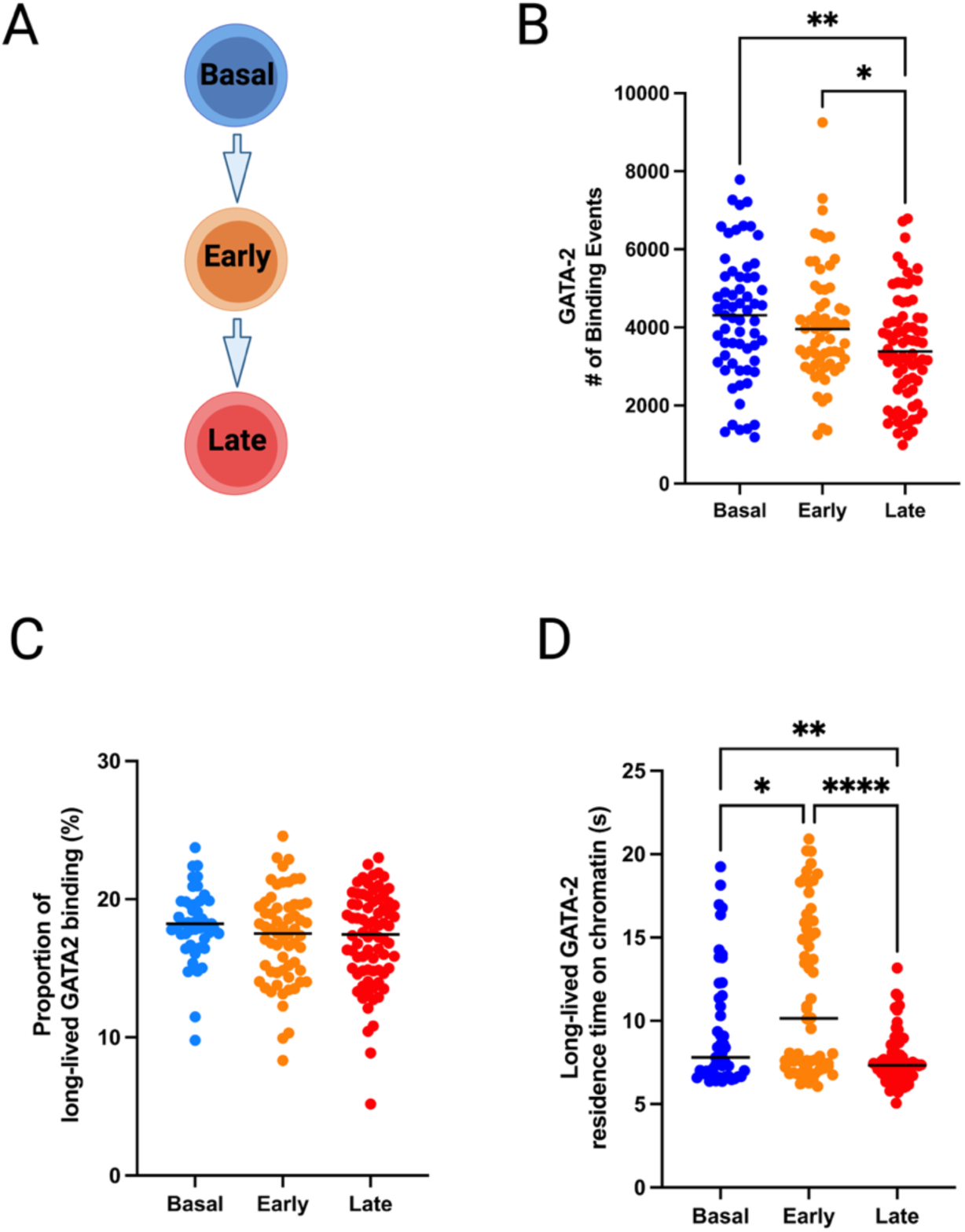
GATA2 binding dynamics exhibit a transitory strengthening of long-lived chromatin engagement during Early erythroid differentiation in G1E-ER4 cells. **(A)** Schematic of the G1E-ER4 differentiation system. Activation of GATA1–ER with 4-hydroxytamoxifen induces progression from the Basal progenitor (0 h) to Early erythroid (2 h) and Late erythroid (24 h) states. **(B)** Total number of detectable GATA2 chromatin binding events per cell across erythroid stages. Binding-event frequency is unchanged between Basal and Early states but is reduced in the Late state relative to both Basal and Early erythroid cells. **(C)** Fraction of GATA2 molecules engaging in long-lived chromatin interactions across erythroid stages. The proportion of long-lived binding events remains constant across Basal, Early, and Late states (approximately 20%). **(D)** Residence times of long-lived GATA2 chromatin interactions across erythroid stages. Residence time is significantly prolonged in the Early erythroid state relative to the Basal progenitor state and is significantly reduced in the Late erythroid state. Each point represents a single cell (Basal, N = 47; Early, N = 62; Late, N = 76). Bars indicate mean ± SEM. Statistical significance was assessed using the Brown–Forsythe and Welch ANOVA with Games–Howell post hoc correction. Significance levels are indicated as follows: * P < 0.05; ** P < 0.01; **** P < 0.0001. Created with BioRender. Hobbs, J. (2026) https://biorender.com/hekoas8

Single-molecule imaging showed that the total number of detectable GATA2 chromatin binding events per cell was unchanged between Basal progenitor and Early erythroid states. In contrast, GATA2 chromatin binding declined by approximately 19% in the Late erythroid state relative to Basal and by approximately 15% relative to the Early state (Figure 2B). Because exogenous GATA2–Halo expression was externally stabilized during imaging, the reduction in detectable binding events in Late erythroid cells likely reflects changes in GATA2’s ability to engage chromatin rather than differences in protein levels. Despite this reduction, the fraction of molecules in the long-lived chromatin binding population remained constant across all stages at approximately 20% (Figure 2C). Early erythroid cells exhibited a significant increase in the chromatin binding residence time of the long-lived binding population relative to Basal progenitors (mean 11.58 vs. 9.54 s; ∼21% increase; p = 0.034). In contrast, residence times declined significantly in Late erythroid cells (mean 7.58 s), representing a ∼21% decrease relative to Basal progenitors (p = 0.0023) and a ∼35% decrease relative to Early erythroid cells (p < 0.0001) (Figure 2D).

Together, these data identify Early erythroid transition as a distinct and transitory kinetic state, characterized by a specific enhancement of GATA2 chromatin engagement dynamics that is not observed in either Basal progenitors or Late erythroid cells. This transitory enhancement occurs at the level of chromatin binding residence time, without expansion of the long-lived binding molecule population. This indicates that Early erythroid entry is marked by changes in chromatin binding behavior rather than increased occupancy. Loss of this kinetic signature in Late erythroid cells is consistent with the established attenuation of GATA2 activity during erythroid commitment, placing this effect within a narrow temporal window of differentiation during which GATA2 chromatin engagement dynamics are transiently enhanced.

### Dynamic regulation of GATA2 chromatin engagement during cytokine mediated HPC7 differentiation

While G1E-ER4 cells have served as a gold-standard model for interrogating the GATA1-induced GATA switch, this system relies on enforced nuclear entry of GATA1 and therefore represents a highly engineered differentiation paradigm. We next asked whether the early strengthening of GATA2 chromatin engagement observed in G1E-ER4 cells also occurs under cytokine-driven differentiation conditions. To address this, we applied a similar single-molecule imaging strategy in HPC7 cells, a hematopoietic progenitor line in which endogenous GATA2 and GATA1 regulation is preserved, and erythroid differentiation is induced by erythropoietin (EPO) stimulation (Supplementary Video 3) (May et al. 2013; Sasca et al. 2019). This approach provides a complementary system in which erythroid entry is initiated by extracellular signaling rather than enforced nuclear localization of GATA1.

Single-molecule imaging in HPC7 cells revealed a pronounced increase in GATA2 chromatin binding events upon cytokine-driven differentiation. The total number of detectable GATA2 chromatin binding events per cell increased significantly from the Basal state to the Early erythroid stage following EPO stimulation and remained elevated in the Late stage (Figure 3B). While the chromatin binding event frequency was markedly higher in both Early and Late erythroid cells compared with Basal progenitors (p < 0.0001), no significant difference was observed between Early and Late stages. To assess functional GATA2 chromatin engagement during cytokine-driven differentiation in HPC7 cells, we quantified the fraction of GATA2 molecules engaged in long-lived chromatin binding events across Basal, Early, and Late erythroid stages. In Basal cells, a mean of approximately 2.6% of total GATA2 binding events were classified as long-lived (>5 s). This long-lived chromatin binding fraction increased significantly during the Early erythroid stage, reaching a mean of 4.4% (p = 0.0067), before declining in Late erythroid cells to levels comparable to Basal (∼2.0%; p = 0.51) (Figure 3C). Notably, Early erythroid cells exhibited a pronounced skew toward higher long-lived chromatin binding fractions at the single-cell level compared with both Basal and Late states. Together, these data indicate that a subset of Early erythroid cells acquires an enhanced capacity for long-lived GATA2 chromatin engagement.

**Figure 3.**
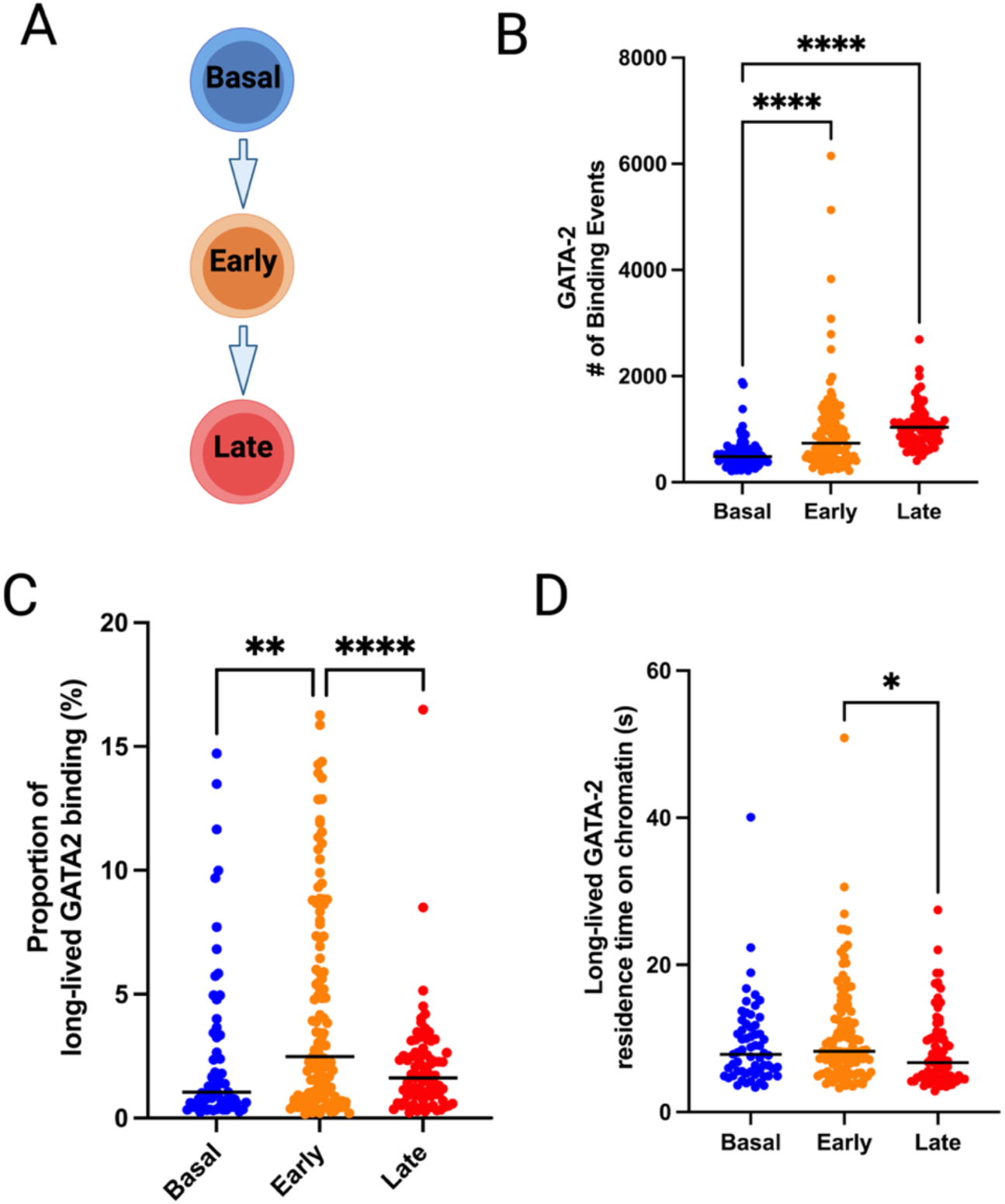
Early erythroid entry in HPC7 progenitors is marked by a transitory increase in long-lived GATA2 chromatin engagement. **(A)** Schematic of the HPC7 erythroid differentiation paradigm. Erythropoietin (EPO) stimulation induces progression from the Basal progenitor state to Early and Late erythroid stages. **(B)** Total number of detectable GATA2 chromatin binding events per cell across erythroid stages. Binding-event frequency increases significantly from the Basal to Early stage following EPO stimulation and remains elevated in the Late stage, with no significant difference between Early and Late populations. **(C)** Fraction of GATA2 molecules engaging in long-lived chromatin interactions across erythroid stages. The long-lived binding fraction is significantly increased in Early erythroid cells relative to Basal progenitors and declines in Late erythroid cells to levels comparable to Basal. **(D)** Residence times of long-lived GATA2 chromatin interactions across erythroid stages. Residence times are similar between Basal and Early stages and are modestly reduced in Late erythroid cells relative to the Early stage. Each point represents a single cell. Bars indicate mean ± SEM. Statistical significance was assessed using the Brown–Forsythe and Welch ANOVA with Games–Howell post hoc correction. Statistical significance was defined as P < 0.05. Significance levels are indicated as follows: * P < 0.05; ** P < 0.01; *** P < 0.001; **** P < 0.0001. Created with BioRender. Hobbs, J. (2026) https://biorender.com/bnwzn4l

We next examined GATA2 chromatin residence times across Basal, Early, and Late erythroid stages in HPC7 cells. The residence time of long-lived GATA2 chromatin interactions was not significantly altered during the Early erythroid transition but was modestly reduced in Late erythroid cells relative to the preceding stage (Figure 3D). This indicates that, in contrast to the Early expansion of the long-lived binding population, changes in GATA2 residence time in HPC7 cells primarily emerge upon the Late stage of erythroid commitment rather than during the initial priming phase.

Short-lived (<1 s) GATA2 interactions were also quantified (Figure Supplement 1C, D). Consistent with the expansion of the long-lived chromatin binding population, the proportion of short-lived binding events decreased in Early erythroid cells. Within the temporal resolution of our measurements, we additionally observed a subtle but significant reduction in the mean residence times of short-lived binding events in Late erythroid cells. Together, these data indicate that the primary characteristics of GATA2 dynamics in HPC7 cells are defined by (i) an Early expansion of the long-lived chromatin binding population during erythroid priming and (ii) a subsequent loss of GATA2 engagement accompanied by reduced residence times of short-lived binding events in the Late stage of erythroid commitment.

These findings indicate that HPC7 progenitors undergo a two-step kinetic transition, characterized by an early expansion of the long-lived binding population during erythroid priming, followed by loss of long-lived GATA2 engagement and reduced residence times upon commitment. Taken together, both examined cell line differentiation models G1E-ER4 and HPC7 converge on a shared principle in which Early erythroid entry represents a transitory kinetic state of enhanced chromatin engagement.

### GATA2 chromatin engagement is transiently enriched in Early erythroid progenitors in vivo

To evaluate GATA2 dynamics under physiological in vivo conditions, we generated a transgenic mouse model in which a SNAP-tag was inserted in frame at the 3′ end of the endogenous *Gata2* coding sequence (GATA2–SNAP; Figure 4A). Prior transgenic mouse models expressing C-terminally tagged GATA2 proteins indicated minimal perturbation of GATA2 function (Ahmed et al., 2020). Accordingly, our SNAP-tagging strategy was designed to preserve endogenous GATA2 expression and regulatory control during hematopoiesis. Consistent with this, the knock-in allele supported normal viability and fertility and did not produce overt hematopoietic abnormalities under the conditions examined, indicating that measurements were obtained in a physiologically competent background.

**Figure 4.**
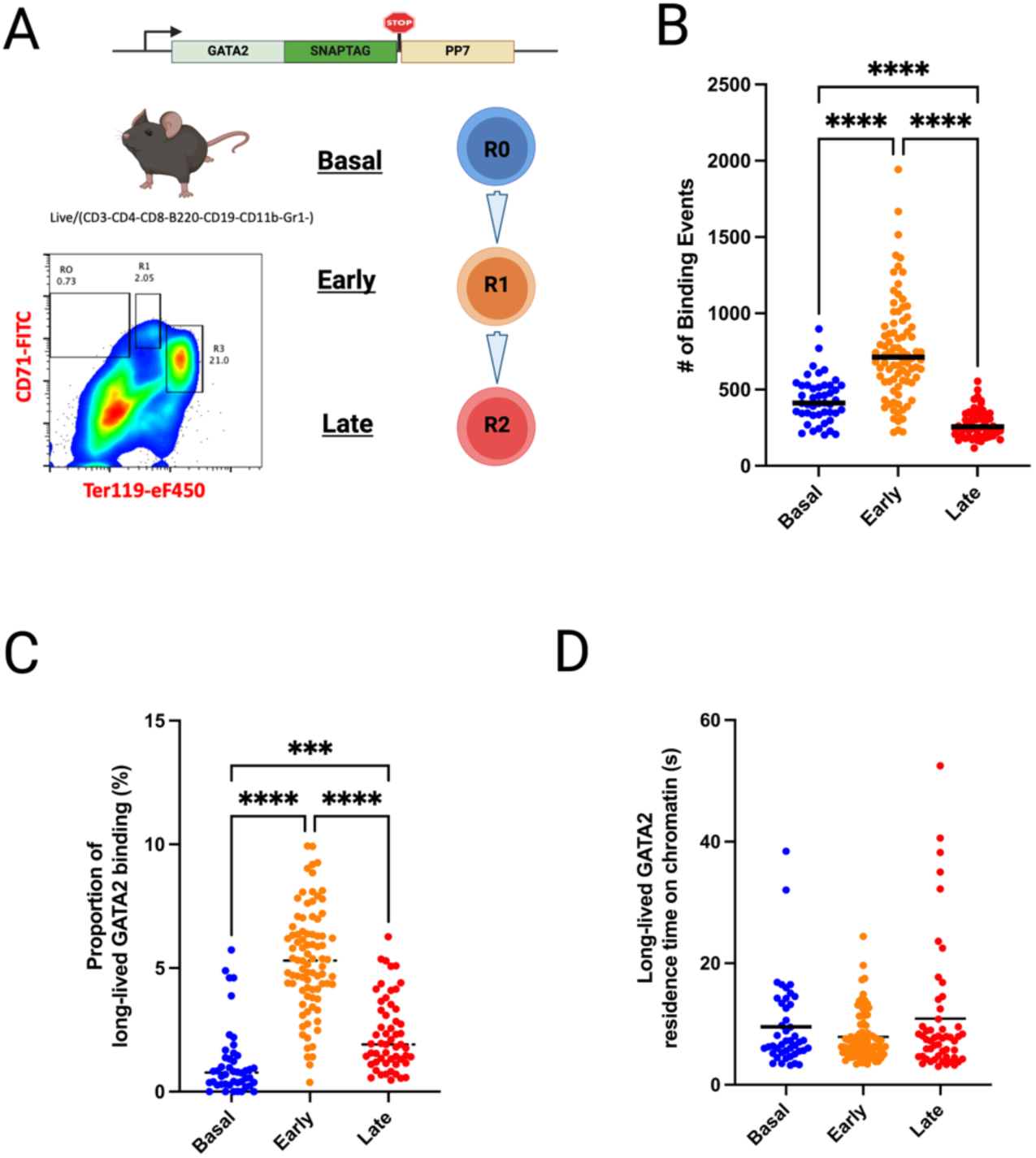
Endogenous GATA2 exhibits transitory enrichment of long-lived chromatin interactions during in vivo erythroid maturation. **(A)** Schematic of the *Gata2*–SNAP knock-in mouse used for endogenous single-molecule imaging, together with the CD71/Ter119 flow-cytometry gating strategy used to define Basal, Early, and Late erythroid populations. **(B)** Total detectable GATA2 chromatin binding events per cell across Basal, Early, and Late erythroid populations. Binding-event frequency is highest in Early cells and reduced in Late cells. **(C)** Fraction of GATA2 molecules engaging in long-lived chromatin interactions across Basal, Early, and Late erythroid populations. The long-lived binding fraction is highest in Early cells and declines upon progression to the Late stage. **(D)** Residence times of long-lived GATA2 chromatin interactions across erythroid stages. Residence times are similar across all three populations, indicating that regulation primarily occurs through changes in the occupancy of the long-lived binding state rather than binding duration. Each point represents a single cell. Bars indicate mean ± SEM. Statistical significance was evaluated using the Brown–Forsythe and Welch ANOVA test with Games–Howell post hoc correction. Statistical significance was defined as P < 0.05. Significance levels are indicated as follows: * P < 0.05; ** P < 0.01; *** P < 0.001; **** P < 0.0001. Created with BioRender. Hobbs, J. (2026) https://biorender.com/ojxcsr1

To define changes in GATA2 chromatin binding dynamics during in vivo erythropoiesis, we isolated primary cells from GATA2–SNAP homozygous mice by flow cytometry to resolve Basal, Early, and Late erythroid populations (Figure 4A). Live-cell single-molecule imaging was performed on these flow-sorted populations by labeling GATA2–SNAP with limiting concentrations of SNAP-Cell 647-SiR SNAP-tag ligand to achieve sparse single-molecule detection, analogous to the HaloTag-based imaging strategy used in the cell line models (Supplementary Video 4). The number of detectable chromatin binding events of endogenously expressed GATA2 increased markedly in Early erythroid cells compared with the Basal population (mean 751.9 vs. 432.3 binding events per cell; ∼74% increase; Figure 4B). In contrast, Late erythroid cells exhibited a substantial reduction in GATA2 chromatin binding events relative to both Basal (mean 276.3; ∼36% decrease) and Early populations (∼63% decrease). This resulted in a three-tier distribution in which Early cells displayed the highest GATA2 chromatin binding event frequency, Basal cells were intermediate, and Late cells showed the lowest levels of chromatin binding (Figure 4B; all pairwise comparisons p < 0.0001). This increase in chromatin binding events in Early erythroid cells may reflect elevated GATA2 protein levels or, alternatively, an increased fraction of GATA2 molecules capable of engaging chromatin.

We therefore quantified the fraction of GATA2 molecules engaging in long-lived chromatin interactions across Basal, Early, and Late erythroid populations. In Basal cells, a mean of approximately 1.2% of GATA2 binding events resulted in long-lived chromatin engagement, whereas this fraction increased markedly to approximately 5.3% in Early erythroid cells (Figure 4C). The fraction of long-lived GATA2 chromatin binding events declined to an intermediate level of approximately 2.3% in Late cells, which remained significantly higher than Basal cells but substantially lower than Early erythroid cells (Figure 4C). These data indicate that Early erythroid cells exhibit a pronounced enhancement in long-lived GATA2 chromatin engagement driven by an expansion of the long-lived binding population, which is partially attenuated as cells progress to the Late stage.

To further define the properties of long-lived chromatin-bound GATA2 molecules, we examined the residence times of long-lived binding events across Basal, Early, and Late erythroid populations. Residence times were similar across all three stages, with no significant differences detected between populations (Figure 4D). Notably, a small subset of multi-second binding events persisted in Late erythroid cells, consistent with continued retention of a limited set of regulatory interactions as transcriptional programs consolidate.

In contrast, short-lived (<1 s) GATA2 interactions exhibited pronounced stage-dependent differences. Residence times increased significantly in Early erythroid cells relative to Basal cells (mean 0.73 s vs. 0.55 s; p < 0.0001) and declined in Late cells (mean 0.58 s), while remaining significantly higher than Basal levels (Figure Supplement 1E–F). Together, these results indicate that long-lived GATA2 engagement is regulated primarily at the level of occupancy of the long-lived binding state rather than its residence time. By contrast, Early erythroid differentiation is accompanied by a slowing of GATA2 target-search dynamics within the short-lived binding population, an effect that is partially reversed later during erythroid differentiation.

In summary, single-molecule measurements in primary cells further support a model in which endogenous GATA2 exhibits a transitory enhancement of long-lived chromatin engagement during Early erythroid differentiation, consistent with the pattern observed in HPC7 cells. While the specific kinetic manifestation slightly differs in the different models, all studied systems including cell lines models and primary cells reveal a conserved Early erythroid window in which GATA2 chromatin engagement is transiently strengthened before it is subsequently attenuated at a later stage of erythroid differentiation.

### Early erythroid–restricted GATA2 peaks resolve promoter-proximal and enhancer-associated regulatory subclasses

Across all systems examined, entry into the early erythroid state is associated with a transitory strengthening of GATA2 chromatin engagement. Leveraging the high degree of synchronization in the G1E-ER4 system to capture this transitory window, we profiled genome-wide GATA2 binding using CUT&Tag to determine whether this kinetic transition is accompanied by specific changes in genomic occupancy. Importantly, chromatin occupancy was mapped using an antibody targeting the HaloTag, leveraging the unique epitope to ensure that detected peaks reflect specific binding of the functional GATA2 fusion protein. Using a consensus peak universe approach, we identified 5,962 GATA2-occupied regions in Basal progenitors, 5,175 in Early erythroid cells, and 4,749 in Late erythroid cells (Figure 5A). Genome-wide CUT&Tag signal heatmaps across each category confirmed stage-restricted enrichment and validated our new temporal classification (Figure Supplement 3).

**Figure 5.**
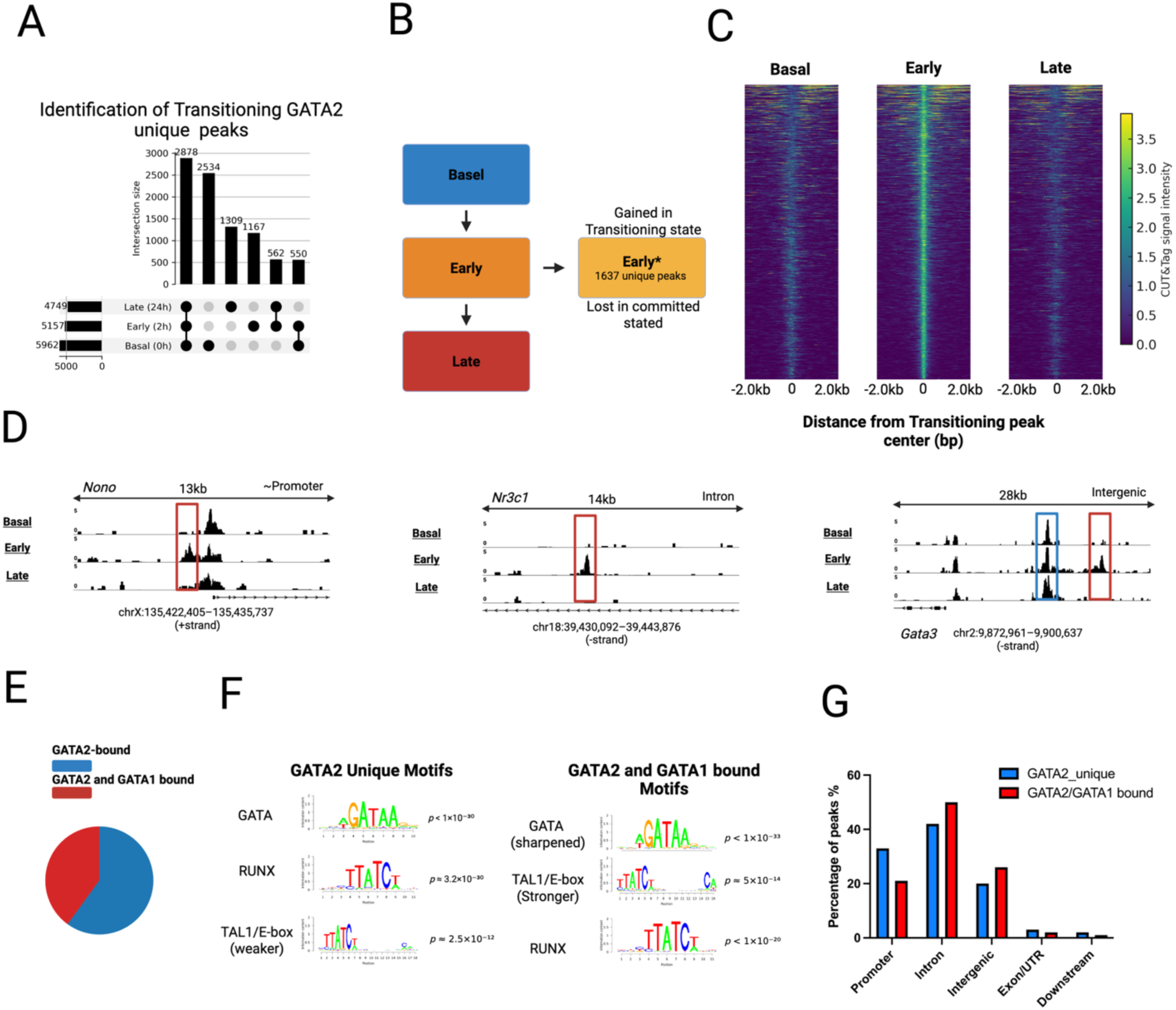
Early erythroid–restricted GATA2 peaks reveal temporally specific recruitment, two regulatory subclasses, and distinct motif and genomic features. **(A)** Categorical classification of GATA2 CUT&Tag peaks across Basal progenitor (0 h), Early erythroid (2 h), and Late erythroid (24 h) G1E-ER4 cells. Peaks were grouped into Early-enriched, shared, Late-acquired, and monotonic categories, providing a global view of how GATA2 genomic occupancy evolves during erythroid differentiation. **(B)** Identification of Early erythroid–restricted GATA2 peaks. A stringent subset of peaks was defined as sites detected by MACS2 exclusively at the Early (2 h) erythroid state (q < 0.01) and absent at both Basal (0 h) and Late (24 h) stages using identical processing parameters. These peaks represent genomic regions uniquely recruited during Early erythroid entry. **(C)** CUT&Tag signal heatmap for all Early erythroid–restricted GATA2 peaks (n = 1,167) aligned to peak centers (±2 kb). Peaks show strong, centered GATA2 enrichment at the Early (2 h) stage with minimal signal at Basal (0 h) and Late (24 h) stages, indicating stage-specific recruitment. **(D)** Genome browser views of representative Early erythroid–restricted GATA2 peaks (highlighted by red boxes), including a promoter-proximal peak near *Nono*, an intronic site within *Nr3c1*, and an intergenic site near *Gata3*. Each locus exhibits GATA2 binding selectively in the Early erythroid state and loss in the Late erythroid state. Blue boxes indicate adjacent genomic regions lacking stage-dependent changes in GATA2 signal. **(E)** Classification of Early erythroid–restricted peaks based on overlap with GATA1 CUT&Tag data. Peaks were subdivided into GATA2-only peaks (n = 700) and peaks bound by both GATA2 and GATA1 (n = 467), indicating that Early GATA2 recruitment can occur either independently of, or in conjunction with, GATA1. **(F)** Motif enrichment analysis for GATA2-only peaks and peaks bound by both GATA2 and GATA1. GATA2-only peaks are enriched for canonical GATA motifs and RUNX-associated elements, whereas peaks bound by both factors display composite GATA/E-box motifs characteristic of TAL1-centered regulatory assemblies. **(G)** Genomic distribution of GATA2-only peaks versus peaks bound by both GATA2 and GATA1 among Early erythroid–restricted sites. GATA2-only peaks are preferentially promoter-proximal, whereas peaks bound by both factors are enriched at intronic and intergenic regions, supporting distinct promoter- and enhancer-associated regulatory roles during Early erythroid differentiation. Created with BioRender. Hobbs, J. (2026) https://biorender.com/u6tblhk

Given the enhanced chromatin engagement observed with live-cell single-molecule imaging in Early erythroid cells, we next focused on isolating genomic regions uniquely occupied during this window. Early erythroid–restricted sites were defined as peaks present exclusively at the Early stage and absent at both Basal and Late stages, yielding 1,167 Early erythroid–restricted peaks (Figure 5B). By requiring complete absence outside the Early erythroid state, this definition yields a narrower, high-confidence peak set that parallels the enhanced chromatin engagement observed in live-cell single-molecule imaging, offering greater specificity than the broader categories identified by intersection analysis.

CUT&Tag signal aligned across all Early erythroid–restricted peaks revealed a sharp, centrally enriched GATA2 signal in Early erythroid cells, with minimal signal at the same genomic coordinates in Basal progenitor or Late erythroid populations (Figure 5C). This pattern, **consistent** with the global stage-specific enrichment profiles (Figure Supplement 3), indicates that these sites represent bona fide stage-restricted GATA2 recruitment rather than persistent binding carried across differentiation states. Representative loci illustrate this behavior across distinct genomic contexts, including a promoter-proximal peak near *Nono*, an intronic site within *Nr3c1*, and an intergenic site near *Gata3* (Figure 5D). Across these diverse genomic contexts, spanning promoter-proximal, intronic, and intergenic regions, each locus exhibits a defined GATA2 peak selectively in the Early erythroid state that is lost again in the Late erythroid state.

To determine whether Early erythroid–restricted GATA2 peaks represent a single regulatory category or multiple subclasses, these sites were intersected with GATA1 CUT&Tag data. This analysis resolved two major groups: GATA2-only peaks (∼60%, n = 700) and peaks bound by both GATA2 and GATA1 (∼40%, n = 467) (Figure 5E). To further characterize these subclasses, we examined their sequence features and genomic annotation. GATA2-only peaks were enriched for canonical GATA motifs and RUNX-associated elements, whereas peaks bound by both GATA2 and GATA1 displayed composite GATA/E-box motifs characteristic of TAL1-centered regulatory assemblies (Figure 5F). Consistent with these motif distinctions, the two subclasses exhibited divergent genomic distributions, with GATA2-only sites showing marked promoter proximity compared to the distal profile of co-bound peaks (Figure 5G).

Together, these analyses indicate that Early erythroid–restricted GATA2 recruitment comprises at least two genomic subclasses defined by transcription factor overlap, motif composition, and genomic annotation. These binding patterns delineate the genomic landscape occupied by GATA2 during Early erythroid differentiation, coinciding with the transitory stage of enhanced chromatin engagement observed by live-cell single-molecule imaging.

## Discussion

The findings presented here demonstrate that GATA2 chromatin engagement is dynamically regulated during erythroid differentiation. Rather than decreasing monotonically as GATA1 levels rise, GATA2 engagement exhibits a distinct temporal reorganization, defining an Early erythroid transition state. Across three complementary systems including cell line models as well as primary cells, live-cell single-molecule imaging reveals a transitory strengthening of GATA2 chromatin engagement during erythroid entry. This strengthening is reflected by prolonged apparent residence times in G1E-ER4 cells under matched acquisition conditions and by an expanded long-lived binding population in HPC7 cells and primary bone marrow progenitors. At the late stage of erythroid commitment, GATA2 chromatin engagement declines across all systems. Together, these coordinated kinetic shifts support a model in which transcription factor binding characteristics are dynamic and not unidirectional, following a non-monotonic trajectory that is closely associated with lineage differentiation.

Prior single-molecule studies have established that transcription factors exhibit both short-lived chromatin interactions (<1 s), typically associated with non-specific target search, and longer-lived interactions (multi-second), which correlate with sequence-specific engagement at regulatory elements (Gebhardt et al. 2013; Hansen et al. 2017; Kenworthy et al. 2022). Interpreted within this framework, the expansion of the long-lived GATA2 binding population and prolongation of residence times observed here indicate enhanced specific chromatin engagement during Early erythroid differentiation. Conversely, the predominance of short-lived interactions in progenitor or late stages is consistent with reduced stable regulatory occupancy at lineage-specific elements.

This kinetic perspective adds a further temporal dimension to the classical GATA-switch model, which has been defined primarily by unidirectional changes in expression and genomic occupancy as GATA1 replaces GATA2 (Bresnick et al. 2012; Suzuki et al. 2013). Our single-molecule data identify a discrete phase at the onset of differentiation during which GATA2 chromatin engagement is transiently intensified before displacement. CUT&Tag profiling supports this framework by identifying regulatory elements occupied by GATA2 only during the Early transition state. These sites segregate into two subclasses: promoter-proximal regions enriched for strong GATA motifs and RUNX1 elements, and distal regions containing composite GATA/E-box motifs characteristic of TAL1-centered enhancer assemblies (May et al. 2013; Tijssen et al. 2011). These subclass-specific features align with known hematopoietic regulatory modules and provide a mechanistic framework for the selective strengthening of GATA2 chromatin engagement at the onset of differentiation.

In this framework, the Early transition state represents a brief phase during which GATA2 increases its engagement with selected regulatory elements before being replaced by GATA1. This model is summarized in Figure 6, which illustrates how GATA2 chromatin engagement is transiently strengthened during early differentiation despite declining protein levels, defining a distinct kinetic phase of the GATA switch. These temporal changes suggest that GATA2 helps establish chromatin configurations that facilitate the subsequent GATA1-driven transcriptional program.

**Figure 6.**
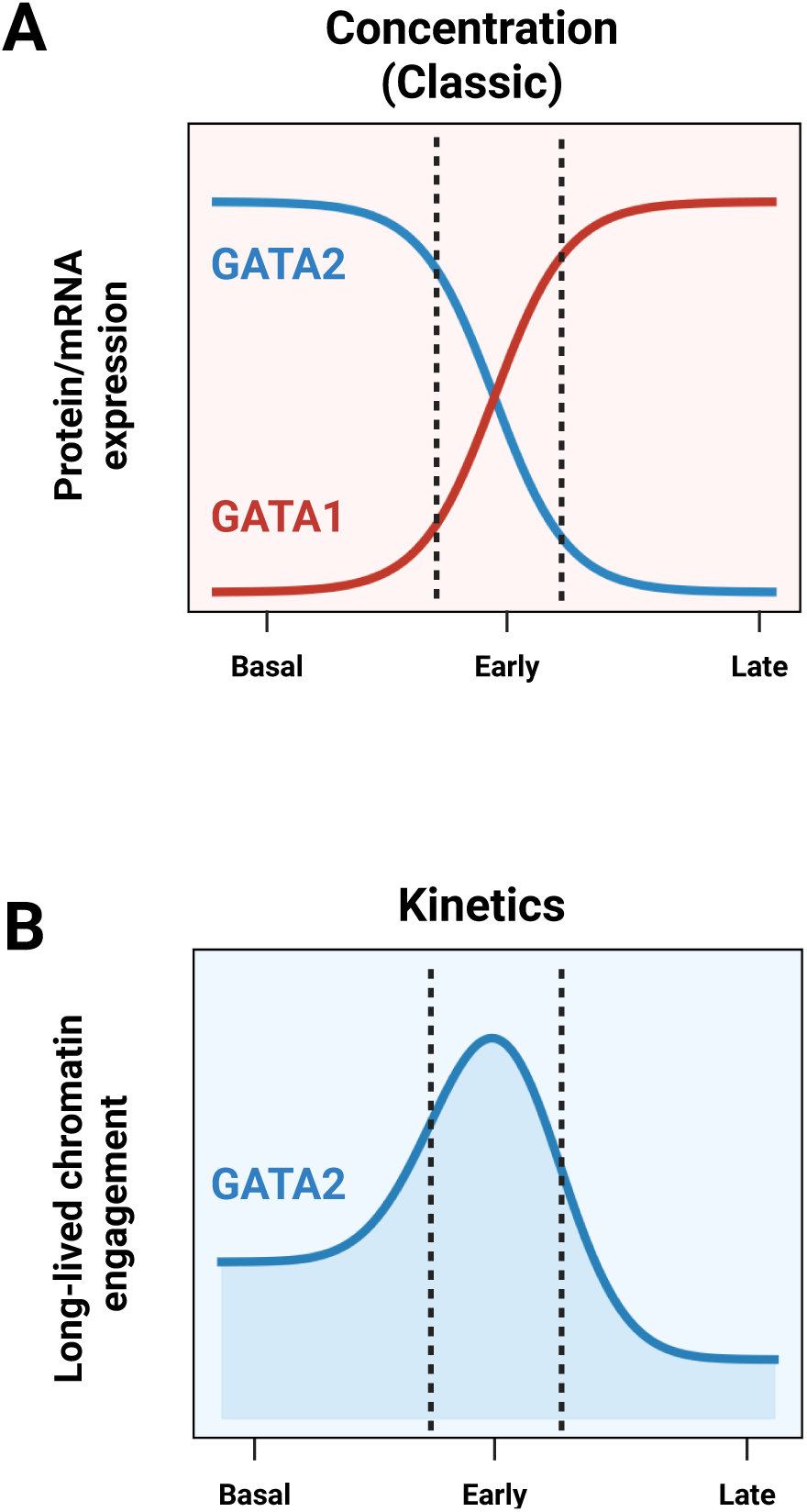
A kinetic transition state uncouples GATA2 chromatin engagement from protein abundance during erythroid differentiation. **(A) Expression Dynamics (Classical View).** The canonical model of the GATA switch, where GATA2 protein levels (blue line) decline monotonically as GATA1 levels (red line) rise to drive differentiation. **(B) Chromatin Engagement Dynamics (Kinetic View).** In contrast to expression, single-molecule measurements reveal that GATA2 chromatin engagement (blue line) is transiently strengthened during the Early erythroid transition. This creates a kinetic peak that uncouples binding activity from protein abundance, defining a distinct biophysical phase of lineage commitment. Created with BioRender. Hobbs, J. (2026) https://biorender.com/cj0q1xh

Comparisons across the three experimental systems indicate that the strengthening of GATA2 chromatin engagement during Early erythroid entry is context dependent. G1E-ER4 cells provide a highly synchronized system in which acute GATA1 activation can be precisely controlled. In this setting, Early erythroid entry is associated with prolonged long-lived GATA2 residence times without detectable expansion of the long-lived chromatin-binding population. The absence of a population-level shift may reflect the engineered GATA1-null background and reduced physiological heterogeneity of this system.

In contrast, HPC7 cells and primary erythroid progenitors exhibit distinct kinetic behavior relative to G1E-ER4 cells. These systems comprise heterogeneous progenitor populations in which cytokine signaling and lineage cues regulate endogenous *Gata2* expression. In both contexts, Early erythroid entry was marked by an expansion of the long-lived GATA2 binding population rather than a pronounced change in residence time. This divergence likely reflects differences in chromatin accessibility, cofactor availability, or transcription factor stoichiometry between engineered and native differentiation systems. Despite these quantitative differences, all three systems share a common qualitative pattern: GATA2 chromatin engagement is transiently strengthened during Early erythroid entry and subsequently reduced in the Late stage as differentiation proceeds.

Primary cells displayed features observed in both cell-line models. As in HPC7 cells, primary Early erythroid progenitors exhibited an increased proportion of long-lived chromatin-bound GATA2 molecules. In addition, primary cells showed an Early-stage increase in the residence time of short-lived GATA2 interactions, followed by a reduction at the Late stage, paralleling the residence-time modulation observed in G1E-ER4 cells. Thus, the single-molecule measurements in primary cells capture both an Early expansion of the long-lived binding population and a coordinated modulation of short-lived interaction kinetics.

Because the knock-in allele is subject to endogenous regulatory control, including native promoter usage and physiological protein stoichiometry, these combined effects likely reflect a more complete regulatory landscape than either engineered cell line systems alone. Notwithstanding, a shared kinetic signature emerges across all models: GATA2 chromatin engagement is strengthened during Early erythroid entry and subsequently diminishes as differentiation proceeds. This convergence supports a unified model in which Early erythroid entry is characterized by a previously unappreciated transitory kinetic phase of enhanced GATA2 chromatin interaction.

Of note, single-molecule imaging was essential for resolving the properties of this Early transition state. While prior single-molecule studies examining mRNA indicate that lineage factors are co-expressed and undergo gradual yet reversible quantitative remodeling during hematopoietic priming (Wheat et al., 2020), population-averaged chromatin binding assays cannot determine whether transcription factor–chromatin interactions follow similarly gradual dynamics. Ensemble approaches, including bulk CUT&Tag, measure chromatin occupancy averaged across large cell populations but cannot capture how transcription factor engagement becomes kinetically strengthened or weakened at the single-molecule and single cell levels. By combining live-cell single-molecule tracking with chromatin profiling, our approach revealed regulatory phases that are not readily apparent from population-averaged measurements alone.

Several methodological advances enabled this analysis: (i) CD43-mediated immobilization of non-adherent hematopoietic progenitors, which preserves native nuclear dynamics in suspension cells; (ii) HaloTag and SNAP-tag labeling strategies, which allow detection of low-abundance transcription factors under sparse-labeling conditions; and (iii) live single-cell single molecule tracking, which captures cell-to-cell heterogeneity in chromatin engagement dynamics. Together, these capabilities establish live-cell single-molecule imaging as a powerful framework for dissecting dynamic transcription factor regulation in hematopoietic differentiation.

The CUT&Tag datasets further suggest that early GATA2 binding may have functional consequences for erythroid lineage progression. Promoter-proximal sites uniquely bound by GATA2 may contribute to maintenance of progenitor identity or priming of early erythroid genes, whereas distal elements bound by both GATA2 and GATA1 may reflect regulatory assemblies that facilitate recruitment of TAL1, LMO2, and additional cofactors during commitment. Cooperative interactions among GATA factors, RUNX1, TAL1, and related hematopoietic regulators have been described in multipotent progenitors and early hematopoiesis (Pimanda et al. 2007; Tijssen et al. 2011). In addition, chromatin architectural factors such as the cohesin complex have been implicated in shaping transcription factor occupancy and lineage bias in hematopoietic cells (Tothova et al. 2017; Viny et al. 2019). Together, these observations raise the possibility that the increased GATA2 engagement observed during Early erythroid entry reinforces specific regulatory networks prior to the subsequent dominance of GATA1, providing a plausible mechanistic basis for the transitory kinetic strengthening depicted in our model (Figure 6).

Although the percentage of long-lived GATA2 chromatin engagement falls during commitment, a minority of long-lived interactions persists. These residual events may help sustain enhancer architecture or maintain limited regulatory output, consistent with GATA-factor roles in enhancer assembly and cis-regulatory organization (Bresnick et al. 2012; Oksuz et al. 2023). Determining whether the remaining long-lived interactions in the Late stage erythroid cells help preserve structural features, reinforce silencing, or maintain minimal regulatory functions will be an important direction for future work.

While our study defines a kinetic framework for the GATA switch, several important limitations should be acknowledged. First, GATA2 functions within multi-subunit regulatory complexes, and tracking GATA2 alone may underrepresent the full heterogeneity of chromatin-binding behaviors exhibited by these assemblies. Second, although CUT&Tag profiling identifies stage-specific genomic regions occupied by GATA2, current live-cell single-molecule imaging approaches cannot yet resolve which specific genomic locus corresponds to an individual binding event *in vivo*. This limitation restricts our ability to directly link changes in binding kinetics to specific regulatory elements within single cells during early erythroid differentiation. Third, although we rely on established literature demonstrating declining GATA2 expression during erythroid differentiation, direct quantification of endogenous GATA2 protein abundance in parallel with kinetic measurements will be an important direction for future work. Finally, our interpretation of the relationship between GATA2 and GATA1 relies on population-based staging rather than direct observation of factor exchange at individual chromatin sites. Future studies using simultaneous dual-color single-molecule tracking will be required to visualize the real-time interplay between GATA2 and GATA1 on chromatin and to determine whether GATA1 actively displaces GATA2 or occupies sites only after GATA2 dissociation. Extending these approaches to additional partner factors (e.g., FOG1, TAL1) or targeted perturbation models will be essential for dissecting how specific protein–protein and protein–DNA interactions give rise to the kinetic states described here.

Overall, our findings indicate that transitory non-unidirectional transcription factor kinetics represent an underappreciated dimension of the GATA switch. Similar temporal shifts in chromatin-binding behavior may occur during the initiation and resolution of other lineage transitions. More broadly, our data demonstrate that single-molecule measurements reveal regulatory phases inaccessible to population-scale methods, establishing transcription factor kinetics as a distinct and critical dimension of lineage differentiation.

## Methods

### Cell culture and differentiation

G1E-ER4 cells were maintained in Iscove’s Modified Dulbecco’s Medium (IMDM) supplemented with 15% FBS, 1% penicillin–streptomycin, monothioglycerol (MTG; Sigma), Kit-ligand conditioned medium, and erythropoietin (EPO; PeproTech #554597), following standard protocols (Weiss et al., 1997). EPO was present throughout both maintenance and differentiation. Cells were cultured at 37 °C with 5% CO₂ in suspension and maintained between 1×10⁵ and 1×10⁶ cells/mL. Erythroid differentiation was initiated with 100 nM 4-hydroxytamoxifen (Sigma #H6278), and cells were collected at Basal (0 h), Early (2 h), and Late (24 h) time points.

HPC7 cells were maintained in IMDM supplemented with 10% FBS, 1% penicillin–streptomycin, 74.8 µM monothioglycerol, and 100 ng/mL stem cell factor (SCF; PeproTech, 300-07). For differentiation, cells were washed and resuspended in medium containing reduced SCF (20 ng/mL) and 4 U/mL EPO according to Sasca et al. (2019). Populations were collected at Basal (0 h, maintenance conditions), Early (2 h post-induction), and Late (24 h post-induction) time points.

Because neither cell line has an established STR reference profile, identity and stability were confirmed by morphology, differentiation behavior, and expression of lineage-appropriate transcription factors. All lines tested negative for mycoplasma (Lonza MycoAlert).

### Generation of stable GATA2–HaloTag reporter lines

Stable reporter lines expressing GATA2–HaloTag were generated by lentiviral transduction. The *Gata2*–HaloTag coding sequence was cloned into the pLVX vector upstream of an IRES–ZsGreen reporter and destabilization domain (DD) (Takara #632175). Lentivirus was produced in HEK293T cells by cotransfecting pLVX-GATA2–HaloTag with psPAX2 (Addgene #63586) and pMD2.G (Addgene #12259) using Lipofectamine 2000. Viral supernatants were collected at 48 h. G1E-ER4 and HPC7 cells were transduced on RetroNectin-coated plates (20 µg/mL; Takara #T100A). After 48 h, ZsGreen-positive cells were isolated by FACS and expanded as stable populations. Expression level and nuclear localization of GATA2–HaloTag were validated by live-cell fluorescence microscopy.

### Induction of GATA2–HaloTag expression

The DD degron promotes constitutive proteasomal degradation of the fusion protein, necessitating stabilization before imaging. Cells were incubated with 1 µM Shield-1 (Clontech) for 2 h prior to single-molecule acquisition. During the final 20 min of incubation, cells were labeled with 5 nM Janelia Fluor 646 HaloTag Ligand (Promega #GA1120), followed by two washes in warm Leibovitz’s L-15 Medium, no phenol red (Gibco #21083027). Shield-1 was used to stabilize the DD-tagged fusion protein prior to imaging and was not applied as an experimental perturbation.

### Generation of Gata2–SNAP–STOP–PP7v3 knock-in mouse line

*Gata2*–SNAP–STOP–PP7v3 knock-in mice were generated using CRISPR/Cas9 genome editing in C57BL/6 zygotes. A donor cassette encoding the SNAP protein, a termination codon, and PP7v3 was inserted immediately downstream of the endogenous *Gata2* coding sequence. Synonymous mutations (P470P, H471H, P472P, S473S) were incorporated into the donor to prevent Cas9 re-cleavage. Founder animals were crossed to wild-type C57BL/6 mice to confirm germline transmission, and heterozygous Gata2–Snap–STOP–PP7v3/+ mice were validated by Sanger sequencing. Animals were subsequently backcrossed to remove potential off-target or random integrations and intercrossed to generate homozygous mice for experimental analysis. All mouse procedures were approved by the Albert Einstein College of Medicine IACUC.

### Bone marrow isolation and flow cytometry

Bone marrow was isolated from the femurs and tibias of homozygous GATA2–SNAP mice (n = 2; 12–14 weeks; male and female). Red blood cells were removed using ACK lysis buffer. Cells were stained for CD71 (clone R17217; Thermo Fisher Scientific, #17-0711-82) and Ter119 (clone TER-119; Thermo Fisher Scientific, #48-5921-82) to define erythroid subpopulations: CD71^high^Ter119^low (proerythroblasts), CD71^high^Ter119^intermediate (basophilic erythroblasts), and CD71^low^Ter119^high (committed erythroid cells).

Sorted populations were collected directly into a Ham’s F-12–based hematopoietic maintenance medium (Gibco, 11765054) supplemented with 10 mM HEPES, 1× penicillin–streptomycin–glutamine (PSG; Gibco, 10378016), 1× insulin–transferrin–selenium–ethanolamine (ITS-X; Gibco, 51500056), 1 mg/mL polyvinyl alcohol (PVA; Sigma, P8136), 100 ng/mL TPO, and 10 ng/mL SCF, following the short-term HSPC culture conditions described previously (Wilkinson et al. 2020). Cells were incubated overnight in this medium. For single-molecule imaging, cells were labeled with 6 nM SNAP-Cell 647-SiR (NEB #S9102S) for 30 min at 37 °C, followed by two washes in warm L-15 medium. Immediately before live-cell imaging, cells were transferred into L-15 medium supplemented with 100 ng/mL SCF and 10 ng/mL TPO and imaged without further culture.

### Microscope acquisition

Immediately prior to imaging, cells were washed in warm L-15 medium and resuspended in L-15 supplemented with 10% FBS and cytokines appropriate for each specific cell state to maintain signaling pathways during acquisition. For G1E-ER4 experiments, imaging medium was supplemented with EPO, with the addition of 100 nM 4-hydroxytamoxifen for Early and Late time points. For HPC7 experiments, imaging medium contained SCF (100 ng/mL for Basal; 20 ng/mL for Early and Late) and 4 U/mL EPO (for Early and Late).

Single-molecule imaging was performed on a Nikon Eclipse Ti microscope equipped with a 150×/1.49 NA oil objective, controlled through VisiView 6.0.0.36 (Visitron Systems). Janelia Fluor® 646 HaloTag® Ligand (Promega #GA1120) or SNAP-Cell® 647-SiR fluorophores were excited using a 639 nm laser, and fluorescence was collected on a Prime 95B sCMOS camera (final pixel size 73 nm). Movies were acquired at 500 ms per frame for >1,000 frames per cell, and a stage-top incubator which maintained samples at 37 °C throughout imaging (Kenworthy et al. 2022).

All movies were acquired under identical illumination power, camera settings, dye concentrations, and preparation conditions across undifferentiated, transitioning, and committed samples, thereby enabling direct comparison of apparent residence time behavior under matched acquisition conditions.

### Image preprocessing

Single-molecule TIFF files were organized and pre-processed using custom Python scripts, ImageJ/Fiji macros, and MATLAB code. Raw TIFF stacks were processed to generate photobleaching-control and background-subtracted TIFF files. Background subtraction was performed using a rolling-ball algorithm with a radius of 50 pixels, and output files were saved into standardized condition-specific directories.

### Single-particle tracking and analysis

Single-particle tracking was performed using the STRAP (Single-molecule TRacking And Processing) pipeline implemented in MATLAB, as described previously (Haque and Coleman 2025). STRAP incorporates automated implementations of SLIMfast and evalSPT (Sergé et al. 2008; Normanno et al. 2015).

Tracking parameters were matched across all conditions and included a numerical aperture of 1.49, an exposure time of 500 ms, a pixel size of 73 nm, a gap allowance of up to 1.5s, and a maximum diffusion coefficient of 0.05 µm²/s. Single-molecule localizations were identified by Gaussian fitting and linked frame-by-frame to generate trajectories.

For each cell, a two-dimensional projection map was generated from the x–y positions of all detected binding events. Nuclear boundaries were defined from this projection, and events occurring outside the nucleus were excluded from downstream analysis. Residence times were extracted for all trajectories and used to classify binding events as transient (<1 s) or stable (>5 s), as described in the main text.

To correct for photobleaching, nuclear fluorescence decay was measured in each cell to determine a per-cell photobleaching lifetime. Apparent residence-time distributions were corrected using a right-censoring model in which the observed off-rate reflects the sum of chromatin dissociation and photobleaching rates. This approach yielded photobleach-adjusted residence-time estimates under constant excitation conditions.

Detailed usage instructions and example workflows for STRAP are provided in the GitHub repository referenced in the Code availability section.

### CUT&Tag

CUT&Tag was performed essentially as previously described (Kaya-Okur et al. 2019; Taylor et al. 2024) with minor adaptations for murine progenitors. Approximately 1×10⁶ cells were lightly fixed with 2% formaldehyde for 2 min at room temperature, quenched with glycine, and immobilized on ConA magnetic beads activated in 10 mM CaCl₂ and 10 mM MnCl₂. Samples were incubated overnight at 4 °C with primary antibodies against HaloTag (Promega G9211) or GATA1 (Santa Cruz sc-265) in Dig-Wash (0.01% digitonin), followed by a 1 h room-temperature incubation with secondary antibody. After washing, pA-Tn5 was added for 1 h at room temperature, and tagmentation was initiated by adding 10 mM MgCl₂ activation buffer for 60 min at 37 °C. DNA was purified by phenol–chloroform–isoamyl extraction. Indexed libraries were generated with NEBNext HiFi 2× Master Mix and amplified for 13 PCR cycles, followed by 1.2× AMPure cleanup. Libraries were sequenced on an Illumina NextSeq 500 (paired-end, 35 bp). Reads were trimmed and aligned with Bowtie2 using recommended CUT&Tag parameters, and peaks were called with MACS2 (q < 0.01, --nomodel --extsize 147). Peak overlaps and stage-restricted sets were computed using bedtools intersect, and heatmaps were generated with deepTools computeMatrix and plotHeatmap. IgG controls were processed in parallel to quantify background and define thresholds during peak calling.

### Computational analysis of CUT&Tag peak sets

Aligned BAM files were processed with deepTools (v3.5.1) and bedtools (v2.29.0). MACS2 peak calling used --nomodel --extsize 147 -q 0.01. Transitioning-specific peaks were defined as MACS2 peaks present only in the transitioning state and absent at 0 h and 24 h (bedtools intersect -v).

### UpSet-derived regulatory categories

Peak sets from the Basal (0 h), Early (2 h), and Late (24 h) states were compared using bedtools and visualized as an UpSet intersection matrix. Peaks were classified as Basal, Early-only, shared, Early-to-Late, Late-acquired, or monotonic. These categories were used for visualization in Figure 5B.

### Signal heatmaps, co-bound classification, and genomic annotation

Signal heatmaps were generated using deepTools computeMatrix (±2 kb around summits) and plotted with plotHeatmap. GATA2-only vs. GATA2+GATA1 bound peaks were determined by intersecting transitioning-specific GATA2 peaks with GATA1 CUT&Tag peaks. Genomic annotations (promoter, intron, exon, intergenic, downstream) were assigned using ChIPseeker.

### Motif enrichment analysis

Motif analysis used SeqPos (Cistrome). BED files corresponding to transitioning-only, GATA2-only, GATA2+GATA1 bound, and region-specific subsets were supplied as input. SeqPos identified enriched known and de novo motifs, including high-information GATA sites, RUNX motifs, and GATA/E-box composite motifs associated with TAL1-dependent enhancers. Genome browser tracks were visualized in IGV 2.16.2.

### Western blotting

Cells were lysed in RIPA buffer supplemented with HALT inhibitors (Thermo #78428). Protein concentrations were determined by BCA assay (Pierce #23227). Equal protein amounts were resolved by SDS–PAGE and transferred to PVDF. Membranes were blocked in 5% milk and incubated overnight at 4 °C with primary antibodies against GATA1 (1:1000) and β-actin (1:5000; CST #4967). After secondary incubation with IRDye-conjugated antibodies, blots were imaged on a LI-COR Odyssey CLx and quantified in Image Studio.

### Statistical analysis

Single-molecule experiments included at least two biological replicates unless noted. Data are shown as mean ± SEM. Each point represents a single cell. Bars represent mean ± SEM values. Statistical significance was evaluated using the Brown–Forsythe and Welch ANOVA test with Games–Howell post hoc correction. Statistical significance was defined as P < 0.05. Significance levels are indicated as follows: *P < 0.05; **P < 0.01. CUT&Tag analyses were performed in R (v4.3) with Bioconductor tools; peak set comparisons used bedtools.

## Supporting information

Supplementary Video 2. Representative single-molecule tracking of GATA2-Halo in G1E-ER4 cells.

Supplementary Video 3. Representative single-molecule tracking of GATA2-Halo in HPC7 cells.

Supplementary Video 4. Representative single-molecule tracking of GATA2-SNAP in primary mouse bone marrow cells.

## Data and Code availability

## Data Availability

The CUT&Tag sequencing data generated in this study have been deposited in the NCBI Gene Expression Omnibus (GEO) and are accessible under accession number GSE318619. Single-molecule tracking datasets and raw movie files are available from the corresponding authors upon reasonable request.

## Code Availability

Single-molecule imaging data were processed using the STRAP (Single-molecule TRacking And Processing) pipeline (Haque and Coleman 2025). STRAP was used for image preprocessing, single-particle tracking, and downstream trajectory analysis. The STRAP scripts are publicly available under the GNU General Public License v3.0 at https://github.com/codedbynayem. Detailed usage instructions are provided in the repository.

## Author Details

**John W Hobbs**

Department of Cell Biology, Albert Einstein College of Medicine, Bronx, NY, 10461 USA; Gruss Lipper Biophotonics Center, Albert Einstein College of Medicine, Bronx, NY, 10461, USA

**Contribution:** Conceptualization, Methodology, Resources, Investigation, Validation, Data curation, Formal analysis, Visualization, Funding acquisition, Writing- Original Draft, Writing-Review and Editing

**Competing Interests:** No competing interests declared

**Samuel J. Taylor**

Department of Cell Biology, Albert Einstein College of Medicine, Bronx, NY, 10461, USA; Department of Pediatrics | Hematology & Oncology, WashU Medicine, St. Louis, MO 63110, USA* Current Address

**Contribution:** Conceptualization, Investigation, Data curation, Formal analysis, Visualization, Writing- Review and Editing

**Competing Interests:** No competing interests declared

**Rajni Kumari**

Department of Cell Biology, Albert Einstein College of Medicine, Bronx, NY, 10461 USA

**Contribution:** Conceptualization, Resources, Investigation, Formal analysis, Visualization, Writing- Review and Editing

**Competing Interests:** No competing interests declared

**Nayem Haque**

Department of Cell Biology, Albert Einstein College of Medicine, Bronx, NY, 10461 USA; Gruss Lipper Biophotonics Center, Albert Einstein College of Medicine, Bronx, NY, 10461, USA

**Contribution:** Methodology, Software, Writing- Review and Editing

**Competing Interests:** No competing interests declared

**Lou Lou Victor Jagaraja**

Department of Cell Biology, Albert Einstein College of Medicine, Bronx, NY, 10461 USA

**Contribution:** Resources, Investigation

**Competing Interests:** No competing interests declared

**Ulrich Steidl**

Department of Cell Biology, Albert Einstein College of Medicine, Bronx, NY, 10461 USA; Department of Oncology, Albert Einstein College of Medicine, Bronx, NY, 10461, USA; Ruth L. and David S. Gottesman Institute for Stem Cell Research and Regenerative Medicine, Albert Einstein College of Medicine, Bronx, NY, 10461, USA; Montefiore Einstein Comprehensive Cancer Center, Bronx, NY, 10461, USA

**Contribution:** Conceptualization, Resources, Validation, Formal analysis, Visualization, Funding acquisition, Supervision, Writing- Original Draft, Writing- Review and Editing, Project administration

**Competing Interests:** No competing interests declared

**Robert A. Coleman**

Department of Cell Biology, Albert Einstein College of Medicine, Bronx, NY, 10461 USA; Gruss Lipper Biophotonics Center, Albert Einstein College of Medicine, Bronx, NY, 10461, USA; Montefiore Einstein Comprehensive Cancer Center, Bronx, NY, 10461, USA

**Contribution:** Conceptualization, Resources, Validation, Formal analysis, Visualization, Funding acquisition, Supervision, Writing- Original Draft, Writing- Review and Editing, Project administration

**Competing Interests:** No competing interests declared

## Funding

**National Institutes of Health, NIGMS R01GM126045**

Robert A. Coleman

**National Institutes of Health (NCI R35CA253127, NCI P30CA013330)**

Ulrich Steidl

V Foundation **(AST2025-007)**

Ulrich Steidl

**National Science Foundation, (NSF GRFP, FAIN-2437848)**

Nayem Haque

**National Institutes of Health, (NIGMS T32GM007491)**

John W Hobbs

**National Institutes of Health, (NIDDK TL1DK136048)**

John W Hobbs

The Funders had no role in the design, implementation or interpretation of results arising from this study, nor any role in deciding on submission of this work to eLife.

## Acknowledgements

We thank Daqian Sun and Jaime Clark from the Flow Cytometry Core at the Ruth L. and David S. Gottesman Institute for Stem Cell Biology and Regenerative Medicine at Albert Einstein College of Medicine for assistance with cell sorting. We also thank Yongwei Zhang of the Gene Targeting and Transgenic Facility for generating the mouse lines used in this study. The authors would like to acknowledge the input and support of all members of the Steidl and Coleman laboratories. This work was supported by a grant from the National Institutes of Health (NIH), including NIGMS R01GM126045 awarded to R.A.C., and NCI R35CA253127 awarded to U.S., as well as V Foundation grant AST2025-007 (awarded to U.S.). R.K. was supported by a Career Development Fellowship from Blood Cancer United. S.J.T. was supported by a Young Investigator award from the Edward P. Evans Foundation. U.S. holds the Edward P. Evans Endowed Professorship in Myelodysplastic Syndromes at Albert Einstein College of Medicine. The Endowed Professorship was supported by a grant from the Edward P. Evans Foundation. Also, this work was supported by Jane A. and Myles P. Dempsey. This work was also supported by NIH NIGMS training grant T32GM007491 and the New York Consortium for Interdisciplinary Training in Kidney, Urological and Hematological Research (NYC Train KUHR) award TL1DK136048 to J.W.H. Schematic diagrams were created with BioRender.com. Grammarly was used for language editing and proofreading during the preparation of this manuscript.

## Supplement

**Supplementary Video 1. Shield-1–dependent accumulation of GATA2–Halo in live HPC7 cells.**

Live-cell fluorescence imaging of HPC7 cells expressing GATA2–Halo–DD labeled with Janelia Fluor® 646 HaloTag® Ligand (green) and counterstained with SPY555-DNA (blue). The split-screen movie compares cells treated with vehicle control (*Left*) versus 1 µM Shield-1 for 2 hours (*Right*). In the vehicle-treated condition (left), little to no GATA2–Halo fluorescence is detected, consistent with proteasomal degradation of the fusion protein. In contrast, Shield-1 treatment (*Right*) restores robust nuclear accumulation of GATA2–Halo without altering nuclear morphology. Movies represent single optical sections acquired under identical illumination and display settings.

**Supplementary Video 2. Representative single-molecule tracking of GATA2-Halo in G1E-ER4 cells.**

Example single-molecule tracking movie illustrating GATA2 chromatin interactions in G1E-ER4 erythroid progenitor cells. Movie shown at 15 fps.

**Supplementary Video 3. Representative single-molecule tracking of GATA2-Halo in HPC7 cells.**

Example single-molecule tracking movie illustrating GATA2-Halo chromatin interactions in HPC7 cells. Movie shown at 15 fps.

**Supplementary Video 4. Representative single-molecule tracking of GATA2-SNAP in primary mouse bone marrow cells.**

Example single-molecule tracking of GATA2 in primary erythroid progenitors isolated from mouse bone marrow. Movie shown at 15 fps.

**Figure supplement 1.**
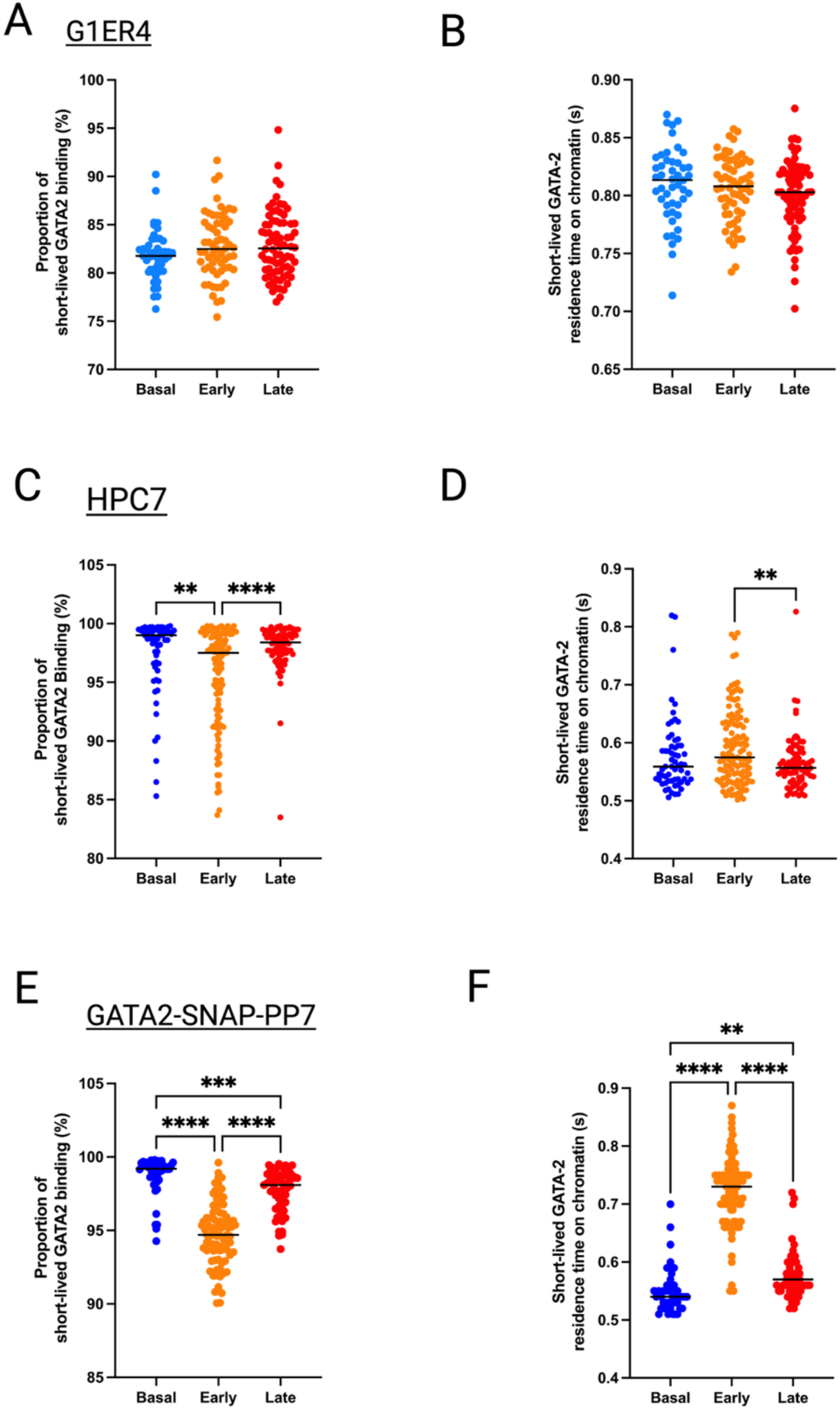
Short-lived GATA2 chromatin binding dynamics across differentiation in G1E-ER4, HPC7, and primary mouse progenitors. **(A–B)** G1E-ER4 cells. The fraction of short-lived (<1 s) GATA2–Halo interactions and the corresponding short-lived residence times did not differ significantly across the Basal (0 h), Early (2 h), and Late (24 h) stages. **(C–D)** HPC7 progenitors. The short-lived fraction decreased modestly at the Early (2 h) stage and increased again at the Late (24 h) stage (C). Short-lived residence times declined significantly from the Early to the Late stage (D). **(E–F)** Primary *Gata2*–SNAP bone-marrow progenitors. The short-lived fraction differed across all three stages, with the largest decrease at the Early stage (E). Short-lived residence times increased from Basal to Early cells and decreased again after the Late stage commitment (F). Statistical comparisons used Brown–Forsythe and Welch ANOVA with Games–Howell post-hoc tests. Significance was defined as P < 0.05, with significance levels indicated as: * P < 0.05; ** P < 0.01; *** P < 0.001; **** P < 0.0001. Created with BioRender. Hobbs, J. (2026) https://biorender.com/r0nnpzm

**Figure supplement 2.**
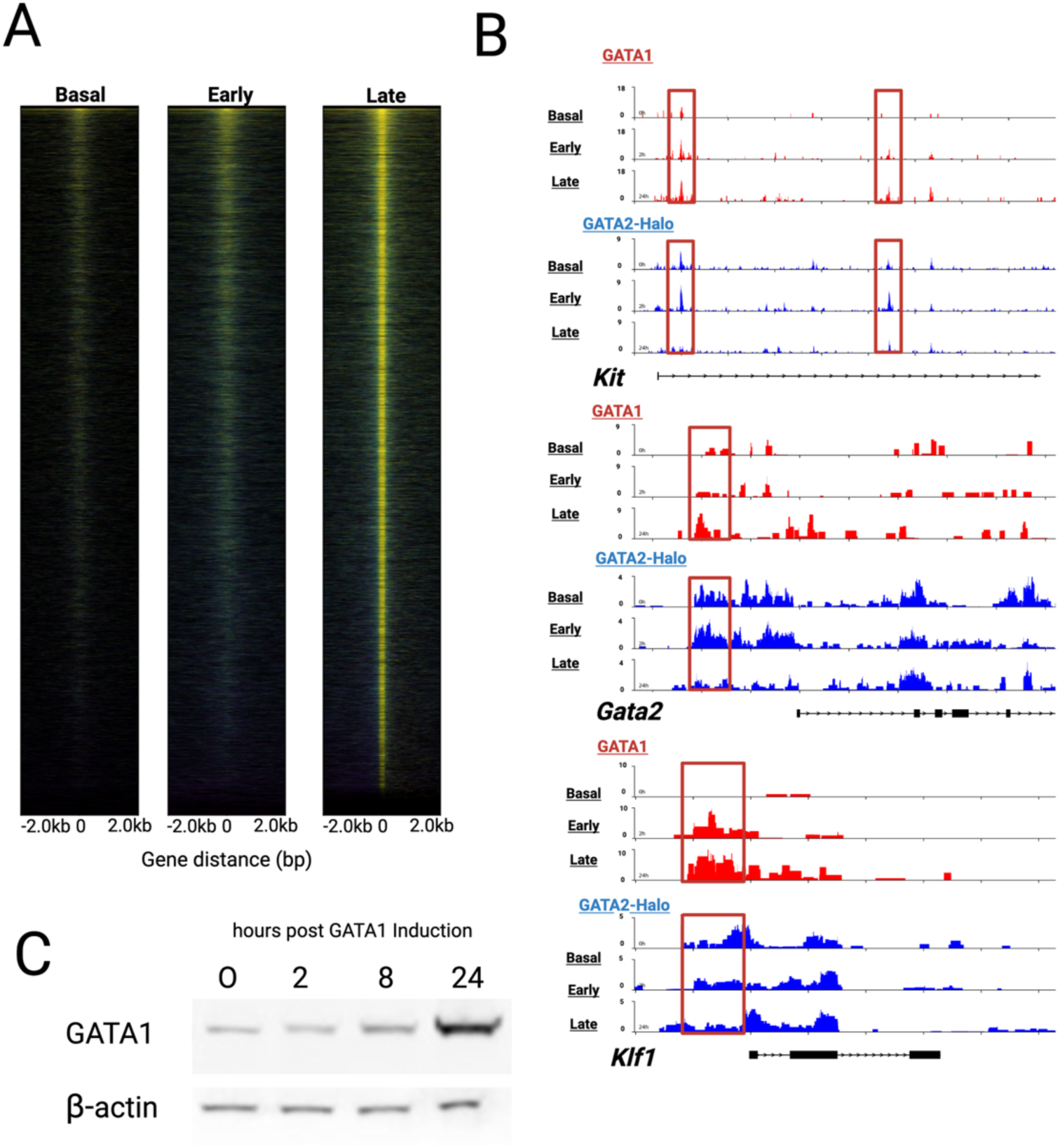
Validation of erythroid induction and GATA factor exchange in the GATA2–Halo reporter line. These analyses confirm that the GATA2–Halo reporter line undergoes normal erythroid induction and GATA1 activation. **(A)** Heatmap of GATA1 CUT&Tag signal aligned across gene bodies at Basal (0 h), Early (2 h), and Late (24 h) stages following 4-hydroxytamoxifen treatment. Rows are ordered by increasing signal intensity. GATA1 occupancy increases progressively over the time course, with strongest enrichment at 24 h. **(B)** Representative genome-browser views comparing GATA1 (red) and GATA2–Halo (blue) CUT&Tag occupancy at the *Gata2*, *Klf1*, and *Kit* loci. Tracks illustrate progressive gain of GATA1 binding accompanied by a reduction or redistribution of GATA2–Halo occupancy during erythroid progression. Red boxes highlight sites of reciprocal GATA factor exchange (“GATA switch”). Y-axis scales are fixed across time points within each factor to enable direct visual comparison. **(C)** Western blot analysis of GATA1 protein abundance at 0, 2, 8, and 24 h after 4-hydroxytamoxifen treatment. β-actin serves as a loading control. GATA1 protein levels increase over time, with maximal expression at 24 h. Created with BioRender. Hobbs, J. (2026) https://biorender.com/weogak3

**Figure supplement 3:**
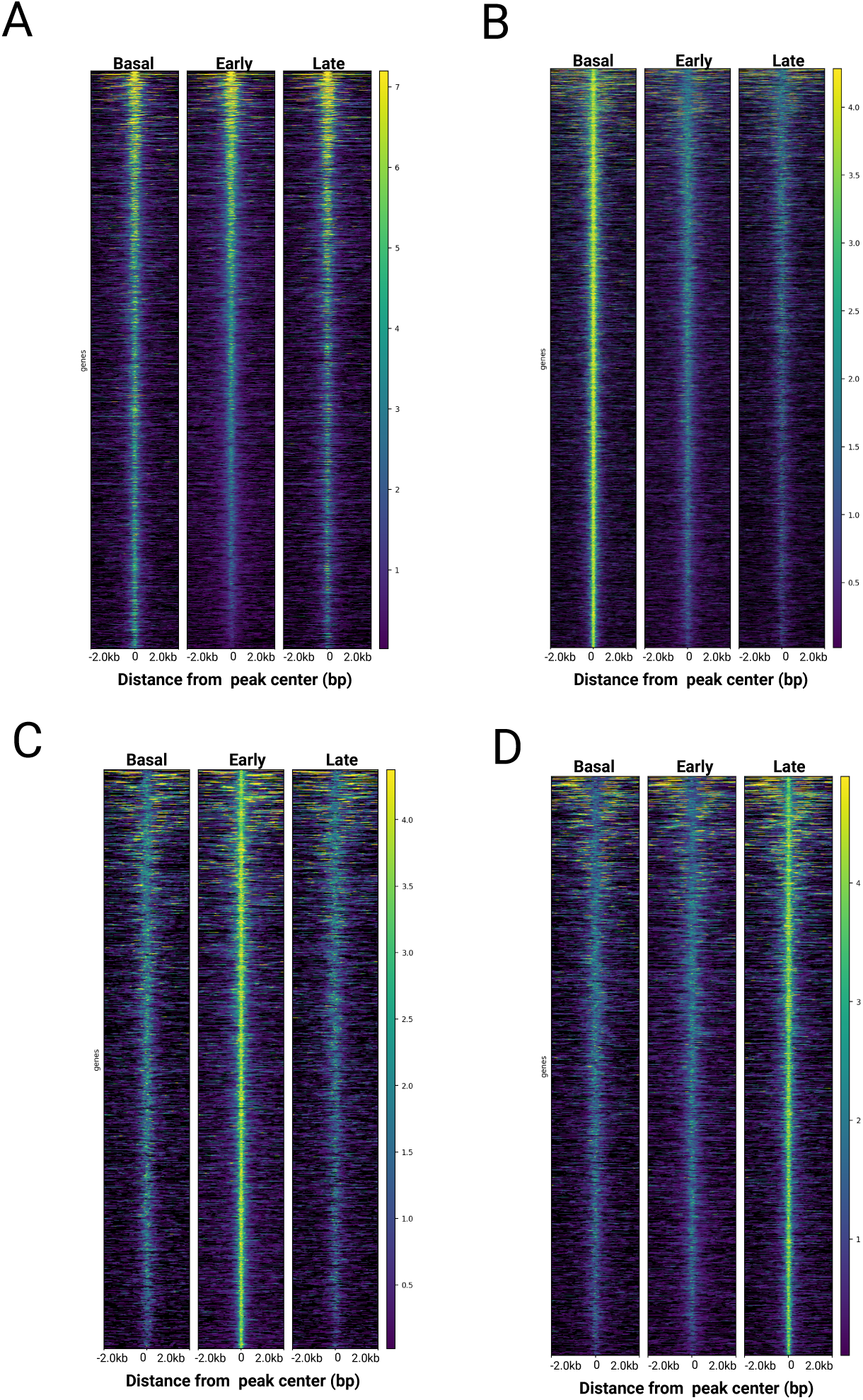
Global CUT&Tag signal patterns across GATA2 peak classes during erythroid differentiation. CUT&Tag signal heatmaps showing GATA2 occupancy across peak categories defined in Figure 5A, including peaks shared across all stages, Basal progenitor–specific (0 h), Early erythroid–specific (2 h), and Late erythroid–specific (24 h) regions. Heatmaps confirm stage-restricted enrichment and validate the temporal classification of GATA2 binding sites used for downstream analyses. **(A)** All GATA2 peaks. **(B)** Basal progenitor–specific peaks (0 h). **(C)** Early erythroid–specific peaks (2 h). **(D)** Late erythroid–specific peaks (24 h). Created with BioRender. Hobbs, J. (2026) https://biorender.com/0tcflll

